# Multiclass Disease Classification from Microbial Whole-Community Metagenomes using Graph Convolutional Neural Networks

**DOI:** 10.1101/726901

**Authors:** Saad Khan, Libusha Kelly

**Affiliations:** Department of Systems & Computational Biology, Bronx, NY, USA; Department of Microbiology & Immunology Albert Einstein College of Medicine, Bronx, NY, USA

**Keywords:** Microbiome, Machine learning, Metagenomics

## Abstract

There is a wealth of information contained within one’s microbiome regarding their physiology and environment, and this is a promising avenue for developing non-invasive diagnostic tools. Here, we utilize 5643 aggregated, annotated whole-community metagenomes from 19 different diseases to implement the first multiclass microbiome disease classifier of this scale. We compared three different machine learning models: random forests, deep neural nets, and a novel graph convolutional architecture which exploits the graph structure of phylogenetic trees as its input. We show that the graph convolutional model outperforms deep neural nets in terms of accuracy (achieving 75% average test-set accuracy), receiver-operator-characteristics (92.1% average AUC), and precision-recall (50% average AUPR). Additionally, the convolutional net’s performance complements that of the random forest, achieving similar accuracy but better receiver-operator-characteristics and lower area under precision-recall. Lastly, we are able to achieve over 90% average top-3 accuracy across all of our models. Together, these results indicate that there are predictive, disease specific signatures across microbiomes which could potentially be used for diagnostic purposes.

## 1. Introduction

There has been an immense interest in the past few years towards developing statistical methods to predict phenotypes such as disease from metagenomic sequencing of one’s microbiome. One of the challenges towards achieving this goal is the problem of separating out signals for different diseases from each other. Unfortunately, many studies that have looked for signatures of individual diseases in the microbiome have produced conflicting results,^1^ and there is evidence that there are general signatures of dysbiosis common to all diseases.^2^ Thus, the standard protocol of comparing samples from a disease of interest against healthy controls, with the goal of identifying features of predictive for that disease, may instead be identifying more general features that signal a diseased or healthy microbiome. This problem can arise if the microbiome signals associated with dysbiosis are stronger than those specific to a given disease. The classifier will then have no mechanism to discriminate between general dysbiosis and the specific signatures of the disease. This lack of discrimination is problematic if we want to understand the differences between diseases on a microbial level and/or make diagnoses for specific diseases. For example, in a clinical setting, a classifier for a certain disease that has not been trained against a diverse range of conditions may produce false positives when applied on a patient who has a different disease.^3^

We propose that approaching this problem as a multiclass classification can alleviate this issue by forcing the classifier to find features in the input that are specific for discriminating between a given class and every other class in the dataset. Additionally, this approach allows us to use a larger dataset containing samples from more conditions, potentially alleviating biases due to batch effects between studies. Unfortunately, making accurate predictions becomes much harder in the multiclass setting, because output is much more specific (random guesses are now correct with probability 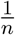 rather than 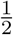). Here, we demonstrate that the multiclass disease classification problem is tractable given the amount of publicly available metagenomic data, we quantify the performance of three machine learning approaches, and we propose graph convolutional network architectures as a powerful method for incorporating microbiome phylogeny for improving performance for deep learning based models. We posit this tractability will only improve over time as more data becomes available.

## 2. Previous Work

Previous studies have used feedforward and convolutional neural networks (CNN) to perform phenotype classifications on smaller datasets than the one considered here, but none have utilized a graph convolutional architecture like the one we describe. Reiman et al. implemented a CNN by embedding phylogenetic trees into ℝ^2^ and used two-dimensional convolutional layers to construct a body-site classifier.^4^ Fioravanti et al. developed a model to diagnose inflammatory bowel disease (IBD) by projecting samples into a two-dimensional space using MultiDimensional Scaling with the patristic distance between phylogenetic trees as the distance metric.^5^ Both papers mapped phylogenetic data to a Euclidean domain to perform convolutions instead of operating in the original tree topology, as we do in this study. Additionally, these studies focused on the disease vs. healthy setting rather than disease vs. all.

More recently, there have been meta-analyses^2,6,7^ that have tried to identify disease-specific signatures which generalize beyond individual studies, but the results presented in these papers have all been for a disease vs. healthy scenario. Our major contribution here is to present three machine learning models, including a novel convolutional architecture which exploits the tree structure of bacterial phylogenies, that can make multiclass disease predictions.

## 3. Problem Setup

There are many moving targets that make machine learning in bioinformatics particularly challenging, and one major problem is the paucity of standardized datasets. Data pre-processing, particularly in the relatively new microbiome field, involves numerous components that are each active areas of research, and thus being continually improved. In the case of the metagenomics data, pre-processing includes new assembly methods, decontamination algorithms, sequencing libraries, annotation methods, reference genomes and so on.^8^ Additionally, new studies are published every month, resulting in an ever-increasing catalog of potential data points to utilize for model training. The result is that different publications compile their own training datasets by aggregating a selection of studies and running a genomic annotation pipeline to pre-process the raw sequence data. This non-standard pre-processing makes comparing models and results between papers difficult because the input datasets can be fundamentally different across each study. Fortunately, there are now actively-maintained standardized datasets to help with this task. For the purpose of this study, we utilized curatedMetagenomicData,^9^ which contains over 7000 whole-community sequenced (WCS) samples across dozens of diseases and multiple body sites. The metagenomic data is represented as a 12365-dimensional vector of taxonomic relative abundances that were generated using the MetaPhlAn2 software package10 (this type of sequencing is also much deeper and higher resolution than traditional 16s rRNA-based sequencing methods).

### 3.1. Dataset Construction

We constructed a dataset containing 5643 samples with 4885 from stool, 403 from skin, 254 from oral cavity, 93 from nares (nasal cavity), and 8 from maternal milk (healthy babies from Asnicar et al.^11^ by including diseases that had at least 15 unique samples. One of the challenges we faced is that some samples have multiple disease labels due to the way the original studies were run. We approached this problem as a multiclasss (one correct label) as opposed to a multilabel (k possible correct labels) problem and thus sought to avoid conflicts due to multiple labeling. Multiple labeling was present in four of our disease sets: Atopic Dermatitis (atopic rhinitis (28), asthma (12)); *C. difficile* (pneumonia (15), cellulitis (2), osteoarthritis (1), ureteral stone (1)); Adenoma (fatty liver (28), hypertension (19), Type II Diabetes (6), Hypercholesterolemia (2), metastases (1)); and Hepatitis B (Schistosoma (1), Hepatitis E (7), Hepatic encephalopathy (2), Hepatitis D (5), Wilson’s disease (1), Cirrhosis (97), Ascites (48)). Additionally, depending on how an individual study was annotated, some samples in the original dataset have no disease label or are instead labeled as “control”. We chose to be conservative in our construction and only included as “healthy” samples where the disease column in the curatedMetagenomicData database was “healthy”.

### 3.2. Graph Convolutional Neural Networks

There are many architectures for generalizing convolutional neural networks to a graph setting,^32^ and we chose to use the method outlined by Kipf and Welling,^33^ which is computationally simple while remaining robust. The method falls into the category of spectral methods, which model the convolution operation as multiplication by a filter operator *g*_*θ*_ in the Fourier domain against input *x* ∈ ℝ^*n*^, where *g*_*θ*_ is a diagonal matrix with parameters *θ* ∈ ℝ^*n*^. The multiplication takes place with respect to the Fourier basis of eigenvectors *U* of the graph Laplacian 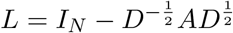, where *D* is the diagonal matrix of vertex degrees, and *A* is the adjacency matrix of the graph (in our case, it is the adjacency matrix of the phylogenetic tree used in our model).

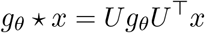

In Euclidean domains, classical convolution operators are equivalent to this operation, and *U* is replaced by the Eigenbasis of the Laplacian operator in ℝ^*n*^, which is the standard Fourier basis {*e*^2*πik*·*x*^: *k* ∈ ℕ^*n*^, *x* ∈ ℝ^*n*^}.

| Disease | Count | Site | Studies |
| --- | --- | --- | --- |
| Atopic Dermatitis (AD) | 38 | Skin | Chng<sup>12</sup> |
| Adenoma | 143 | Stool | Thomas,<sup>6</sup> Feng,<sup>13</sup> Hannigan,<sup>14</sup> Zeller<sup>15</sup> |
| Bronchitis | 18 | Stool | Yassour<sup>16</sup> |
| *C. difficile Infection (CDI)* | 33 | Stool | Vincent<sup>17</sup> |
| Colorectal Cancer (CRC) | 273 | Stool | Vogtmann,<sup>1</sup> Yu,<sup>18</sup> Feng,<sup>13</sup> Zeller,<sup>15</sup> Hannigan<sup>14</sup> |
| Fatty Liver | 94 | Stool | Loomba,<sup>19</sup> Feng<sup>13</sup> |
| Hepatitis B Virus (HBV) | 99 | Stool | Qin<sup>20</sup> (2014) |
| Healthy | 3808 | All | 25 Studies |
| Hypertension | 169 | Stool | Thomas,<sup>6</sup> Feng,<sup>13</sup> Li<sup>21</sup> |
| Inflammatory Bowel Disease (IBD) | 148 | Stool | Nielsen<sup>22</sup> |
| Impaired Glucose Tolerance (IGT) | 49 | Stool | Karlsson<sup>23</sup> |
| Infectious Gastroenteritis | 20 | Stool | David,<sup>24</sup> Yassour<sup>16</sup> |
| Metabolic Syndrome | 50 | Stool | Vrieze<sup>25</sup> |
| Otitis | 107 | Stool | Yassour<sup>16</sup> |
| Periodontitis | 48 | Oral | Shi<sup>26</sup> |
| Psoriasis | 74 | Skin | TettAJ<sup>27</sup> |
| Rheumatoid Arthritis (RA) | 194 | Stool | Chengping<sup>28</sup> |
| Type I Diabetes (T1D) | 55 | Stool | HeintZ-Buschart,<sup>29</sup> Kostic<sup>30</sup> |
| Type II Diabetes (T2D) | 223 | Stool | Qin<sup>31</sup> (2012), Karlsson<sup>23</sup> |

Since there are an infinite number of Fourier modes for any domain, we choose a cutoff K and use the first K Laplacian eigenvectors when constructing *U*. However, this operation is computationally expensive on graphs because there is no general analogue of Fast Fourier Transforms outside of a Euclidean domain (meaning we would have to perform direct matrix multiplications), and the Eigenbases cannot be precomputed because they are different for each graph. The method of Kipf and Welling overcomes this problem by setting *K* = 1 and reformulating the above operation to get

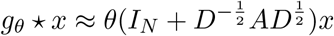

The choice of cutoff specifies that all nodes up to *K* edges away from a given node contribute to the output of the convolution operator at that node. Thus, *K* = 1 implies that the output at each node is a function of that node and its immediate neighbors. By stacking layers of this form together (after applying a non-linearity at each level), we can integrate information from increasingly farther nodes. Each individual convolution layer can thus be written as

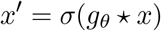

for some non-linearity *σ*. Lastly, this framework can be extended to incorporate more than one input / output channel per layer by adding an additional channel dimension to each of these parameters (see original paper for details).

**Fig. 1.**
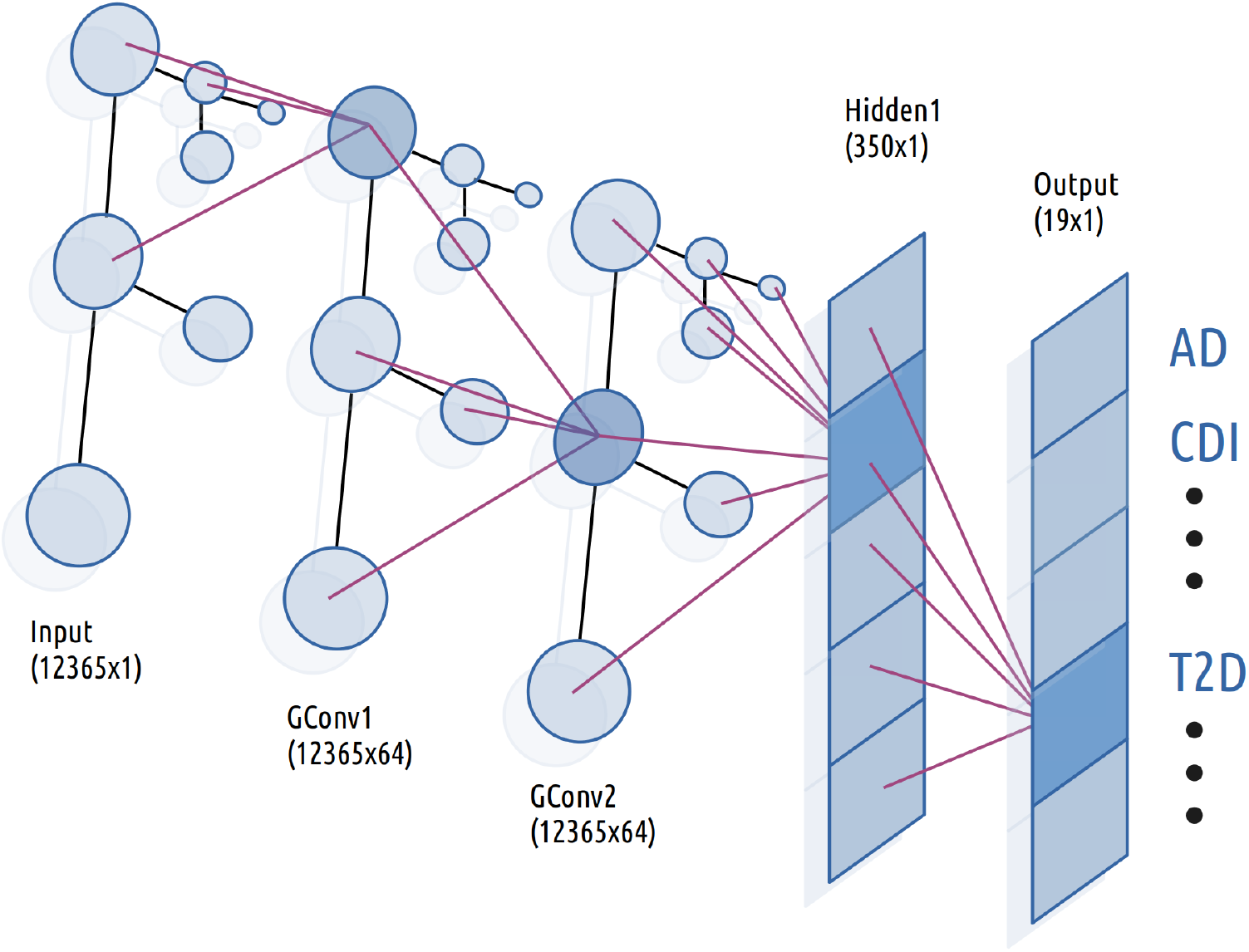
Architecture of Graph Convolutional Classifier. Purple lines indicate the flow of information from the previous to the highlighted neuron in each layer. In the convolutional layers, each neuron receives information only from its immediate neighbors in the preceding layer.

### 3.3. Models

We constructed our graph convolutional neural network (GCN) model by stacking two GCN layers with 64 channels each followed by a fully-connected linear layer with 350 nodes. We used exponential linear units (eLU) as our activations between each layer and a sigmoid activation at the top level for classifications.^34^ Our standard deep neural network (DNN) model consisted of two fully connected linear layers with eLU activation with 1000 and 350 neurons respectively followed by a sigmoid classification layer. Additionally, the GCN model was initialized with the adjacency matrix of the phylogenetic tree corresponding to the taxa present in our 12365-dimensional input vector. All three classifiers took such a 12365-dimensional abundance vector as input. We implemented our neural networks using PyTorch along with the PyTorch Geometric library for the GCN components.^35,36^ Lastly, our random forest model was constructed using the default settings from the `sklearn.ensemble.RandomForestClassifier` module except for the number of trees, which was set to 1000.^37^ We settled on these configurations after manual experimentation. We found that increasing the number of trees improved generalization performance of the random forest up to a certain point, and the size of hidden layers in our networks did not make a dramatic difference in performance.

### 3.4. Training

The biggest challenge in this study was dealing with the extreme class imbalance caused by more than half of the samples in our dataset coming from healthy patients and many diseases having only a few dozen samples. We found that any classifier trained directly on this dataset demonstrated extremely poor generalization performance and/or devolved to classifying every sample as the same label (healthy) regardless of input. We overcame this hurdle by employing a combination of class weighting and oversampling. During each experiment, we constructed an oversampled training set by computing the size of the largest class in the training set (healthy) and for every other class, randomly resampling from that class until each class had the same number of samples.^38^

We augmented resampling by assigning each class a weight of 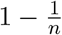, where *n* is the number of classes, and training our neural networks in one-vs-all manner for each class (using a binary cross-entropy loss function).^39^ This approach is equivalent to training *n* different classifiers with shared weights, one for each class, where the objective of each classifier is to discriminate between its class and every other sample in the dataset. To compute the top prediction of the classifier, we then ran the outputs of the network through a softmax function and returned the index of the highest class. This is a commonly used technique in multiclass classification settings, because it reduces the difficult problem of discriminating between *n* classes to *n* easier problems of discriminating between 2 classes.^40^ We used a 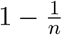 weight in order to address a new class imbalance problem that arises in this setting; if there are *m* examples of each class in the dataset after oversampling, then each binary classifier will see *m* positive examples and *mn* negative examples. A 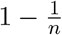 weight magnifies the importance of the positive samples for each class in order to compensate for this. We used a 70-30 split between training and test set, rather than the usual 80-20 in order to have sufficient test samples for each class to evaluate; additionally, we generated a new train-test split for each training run and ran 30 iterations of the GCN model, 30 of the DNN, and 20 of the random forest.

Next, we trained our classifiers. Data preprocessing, ingest, and analysis for our neural networks was performed using Fireworks, a PyTorch-based library that we previously developed to facilitate common machine learning tasks.^41^ The GCN and DNN were trained using the Adam optimizer with 2 ∗ 10^−5^ and 1 ∗ 10^−^ respective learning rates, 40 and 100 respective batch sizes, weight decay parameter set to 1, binary cross-entropy loss, and an early stopping condition when the loss dropped to 2. We performed an analogous procedure with our random forest classifier by assigning the same class weights and using the same over-sampled training set as with the neural network, and we trained the model using its `.fit` method.

## 4. Results

The average accuracy varied widely by disease and classifier, indicating the difficulty of this problem given the number of positive samples for each category. However, most of the the classifications were statistically better than random (5.3%). In general, the convolutional net performed on par with or better than the deep net in terms of accuracy, and both models had higher accuracy than the random forest for most diseases except for AD, HBV, RA, and hypertension. The random forest excelled in the healthy vs. disease task, producing an impressive 99% accuracy. Because healthy was also the largest component of the test set, constituting 66.9% of samples on average, this was responsible for the bulk of its performance in terms of weighted accuracy. However, overall accuracy can be biased by the distribution of the test set (ie. if there were more samples for the classes that a given classifier excelled at, then that classifier would benefit). When we weighted each class equally, then the GCN had the highest average accuracy (46% vs 34.6% for random forest and 44.9% for DNN), indicating that it was more accurate across a broader range of diseases than the random forest model.

Additionally, we often want the ability to tune our models in order to have a trade off between sensitivity and specificity. The AUC, or the area under the receiver operating characteristic curve (ROC), is one such measurement which is invariant to label distributions. We generated ROCs for our neural nets by varying the bias threshold of the final layer and evaluating their true positive and false positive rate on the test set, and for the random forest by using the `sklearn.metrics.roc_curve` function on the test set predictions. We found that the GCN model had higher or statistically equivalent AUCs across all labels than both the random forest and the deep net. In particular, we achieved an average AUC of 89.5% for T2D, which previous studies have found to be a particularly challenging task^7^ (AUCs typically range in the 60s).

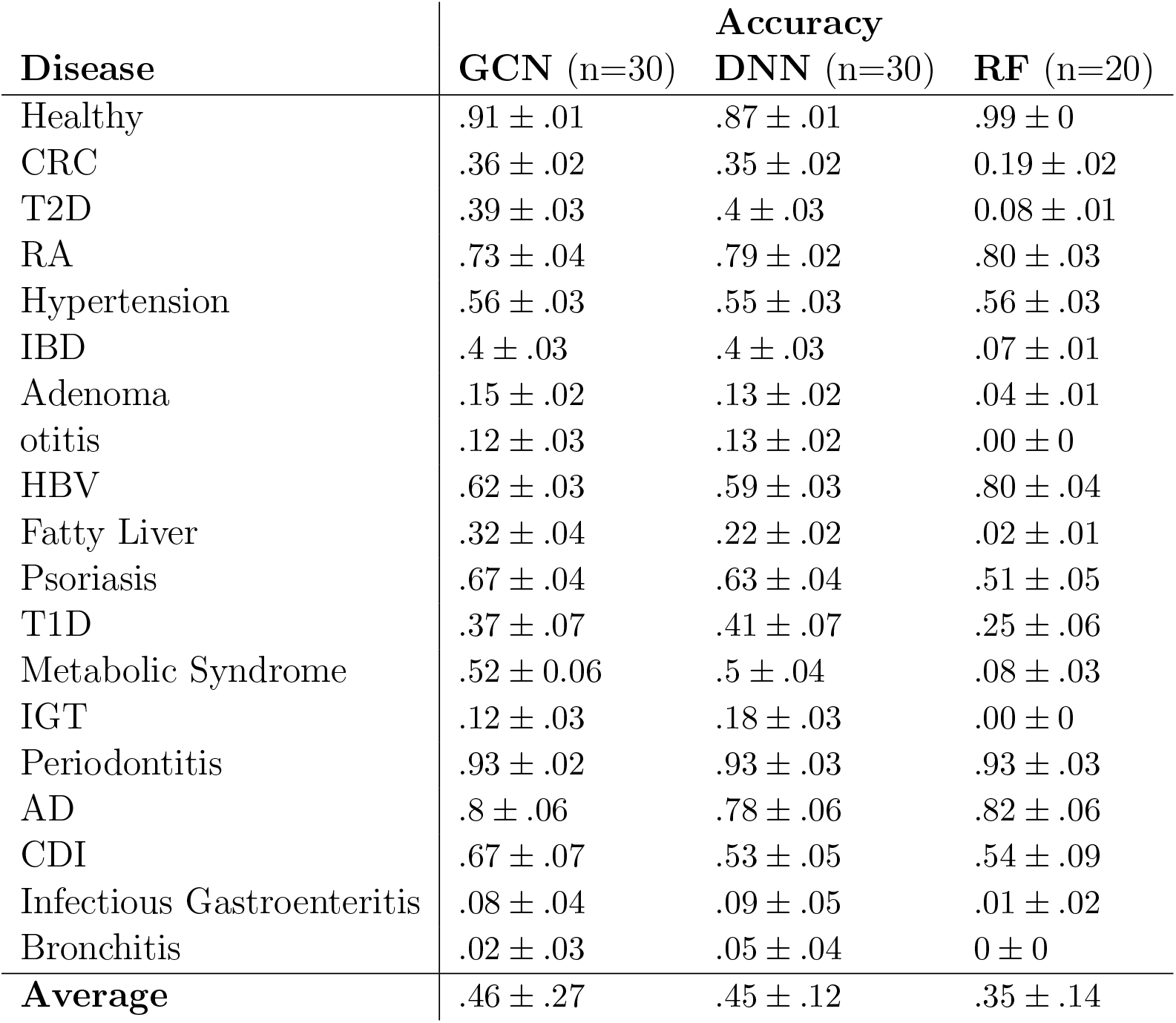

While problematic, class-imbalanced datasets are also the norm in reality, especially if we are interested in the clinical context where most patients do not have the disease being screened for. While ROC curves are a useful class-distribution invariant performance metric, they can also mask the effect of a large false-positive rate. We thus computed precision-recall curves (PRC), which plot the relationship between positive-predictive-value (precision) and sensitivity (true-positive rate), in order to compare these classifiers in the context of class imbalance. We found that the random forest performed much better than the two neural network models with respect to the area under the PRC, implying that its predictions are more reliable in the context of class imbalance.

| Disease | AUC |  |  | AUPR |  |  |
| --- | --- | --- | --- | --- | --- | --- |
|  | GCN | DNN | RF | GCN | DNN | RF |
| Healthy | .84 ± .01 | .82 ± .0 | .83 ± 0 | .73 ± .08 | .88 ± .01 | .93 ± 0 |
| CRC | .84 ± .01 | .81 ± .02 | .5 ± 0 | .41 ± .04 | .35 ± .03 | .53 ± 0 |
| T2D | .9 ± .01 | .86 ± .01 | .5 ± 0 | .4 ± .05 | .31 ± .02 | .52 ± 0 |
| RA | .98 ± .01 | .99 ± .0 | .85 ± .02 | .73 ± .04 | .74 ± .03 | .85 ± .02 |
| Hypertension | .94 ± .01 | .92 ± .01 | .66 ± .01 | .51 ± .05 | .44 ± .03 | .6 ± .02 |
| IBD | .94 ± .02 | .9 ± .01 | .5 ± 0 | .41 ± .04 | .36 ± .02 | .52 ± 0 |
| Adenoma | .81 ± .02 | .75 ± .02 | .5 ± 0 | .15 ± .03 | .12 ± .01 | .51 ± 0 |
| Otitis | .86 ± .02 | .74 ± .02 | .5 ± 0 | .18 ± .03 | .1 ± .01 | .51 ± 0 |
| HBV | .97 ± .01 | .96 ± .01 | .69 ± .02 | .64 ± .05 | .58 ± .03 | .7 ± .02 |
| Fatty Liver | .89 ± .02 | .8 ± .02 | .5 ± 0 | .35 ± .07 | .2 ± .03 | .51 ± 0 |
| Psoriasis | .98 ± .01 | .93 ± .02 | .73 ± .03 | .74 ± .03 | .57 ± .04 | .62 ± .03 |
| T1D | .98 ± 0 | .97 ± .01 | .53 ± .03 | .47 ± .07 | .48 ± .07 | .53 ± .02 |
| Metabolic Syndrome | .98 ± .01 | .97 ± .01 | .51 ± .01 | .63 ± .08 | .55 ± .03 | .51 ± .01 |
| IGT | .95 ± .01 | .88 ± .02 | .5 ± 0 | .22 ± .03 | .16 ± .01 | .5 ± 0 |
| Periodontitis | .99 ± 0 | .99 ± 0 | .96 ± .02 | .95 ± .02 | .93 ± .02 | .96 ± .02 |
| AD | .99 ± 0 | .99 ± 0 | .88 ± .03 | .89 ± .04 | .78 ± .05 | .85 ± .03 |
| CDI | .99 ± 0 | .91 ± .04 | .64 ± .03 | .75 ± .04 | .64 ± .08 | .62 ± .03 |
| Infectious Gastroenteritis | .88 ± .03 | .7 ± .1 | .5 ± 0 | .21 ± .04 | .14 ± .01 | .5 ± 0 |
| Bronchitis | .78 ± .04 | .74 ± .06 | .5 ± 0 | .13 ± .03 | .13 ± .02 | .5 ± .0 |
| **Average** | .92 ± .03 | .88 ± .12 | .62 ± .07 | .5 ± .11 | .45 ± .12 | .62 ± .07 |

Next, we analyzed the ranking of predictions by our models. In a clinical setting, a ranked ordering of probabilities may be more useful than a single output, because a physician can integrate that information along with the rest of their resources to generate a differential diagnosis. We examined the accuracy of the top-3 and top-5 predictions that each classifier produced. Most diseases were correctly identified within the top-3 classifications, and almost almost every disease was correctly present with at least 90% of the time in the top-5 classifications for each model. We measured which classes were most often predicted for a given label when an incorrect prediction was made for a subset of the labels (ie. which how often the model confused a given disease for another disease). Healthy was often misclassified as T2D or CRC, and every disease was often misclassified as healthy. These patterns were also consistent across the three models. These errors may simply be due to the limited size of the dataset or noise in the system. But it could also indicate that the concept of a ‘healthy microbiome’ is vacuous given the broad of range of microbiomes that a healthy individual can have. Additionally, diseases can manifest with a broad range of severities, which may result in different metagenomic signatures which would all get grouped under a single label.

**Fig. 2.**
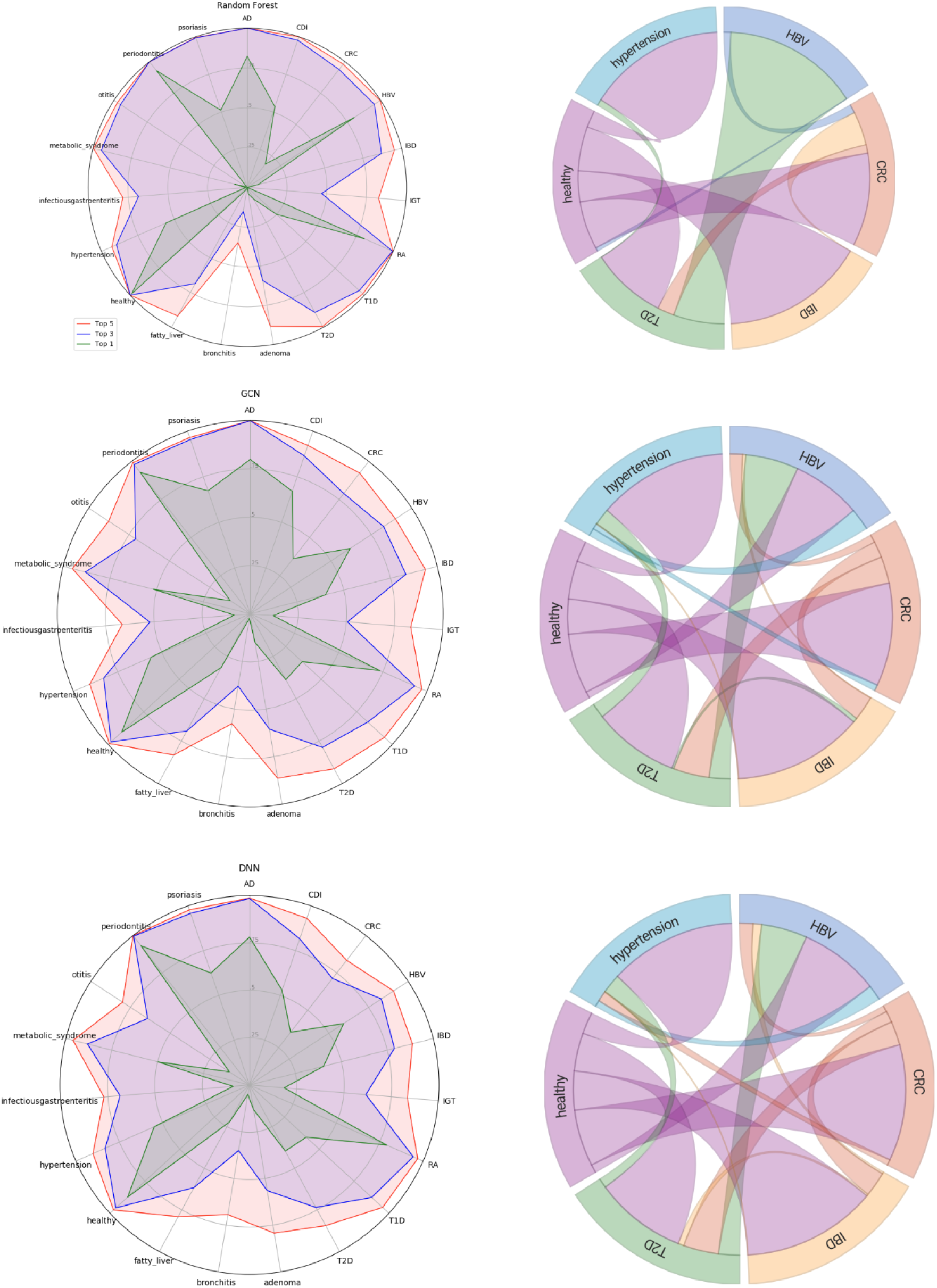
(left) Accuracy at top-1,3, and 5 levels for (top to bottom) Random Forest, GCN, DNN. (right) Chord diagram showing (for a subset of labels) the most common classification made when an incorrect classification was made for a given class.

**Fig. 3.**
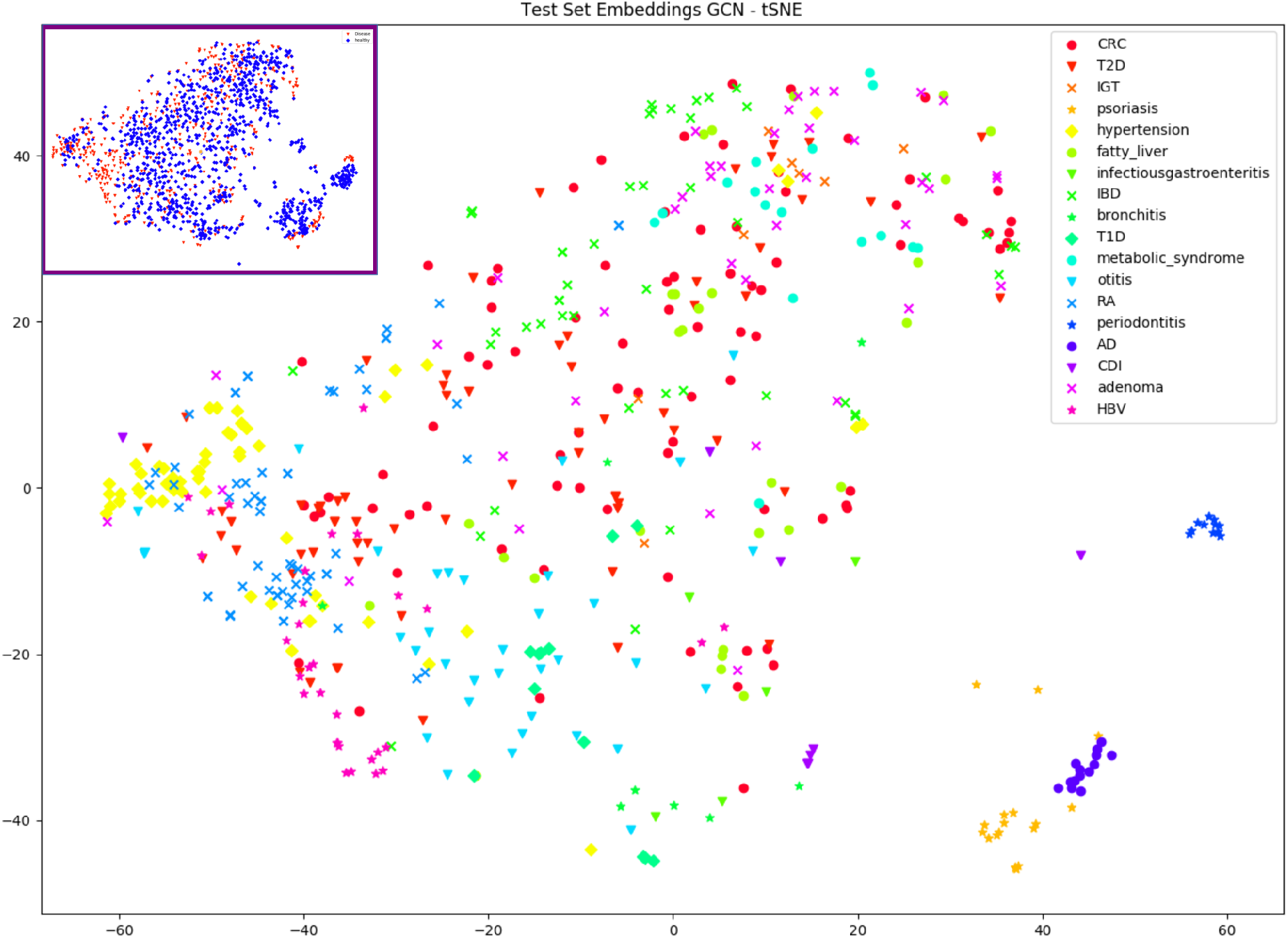
tSNE visualization of the final layer activations of the GCN model (excluding healthy). Inset shows the same tSNE labeled by healthy (blue) and disease (red).

Lastly, we visualized the hidden layer activations of our GCN model on the test set using tSNE to see how different diseases cluster. We found that the skin conditions AD and psoriasis clustered together, separate from the other diseases which were evaluated on stool samples. Periodontitis, which was represented via oral microbiome samples, also clustered independently. Some other conditions, such as hypertension, adenoma, and otitis weakly clustered in some regions of the graph. We also visualized healthy and disease using the same tSNE, and we found that healthy microbiomes were scattered throughout the plot alongside disease, consistent with the idea that there is a broad range of possible healthy microbiomes.

## 5. Conclusion

We have extended the results of previous work on microbiome-phenotype prediction here by demonstrating that multiclass disease prediction from whole community metagenomes, a clinically relevant task for machine learning, is a tractable problem and is improved by using the taxonomic structure of bacterial communities. We implemented multiple classifiers that were able to discriminate between 18 different diseases and healthy with greater than 70% accuracy on a dataset of over 7000 samples. Moreover, while the GCN model generally outperformed the DNN, the RF excelled on a completely different set of metrics, indicating that these two models could potentially complement one another. RF based models, achieving 99% accuracy on healthy vs. not-healthy, could be used to identify dysbiosis in general, while GCNs could potentially then be used to discriminate between individual diseases. Additionally, the success of the GCN model implies that the geometric structure entailed in microbial phylogenies contains meaningful information for disease classification. This is particularly exciting result because graph and tree data structures are ubiquitous in systems biology, so GCN architectures may be applicable to a broad range of problems. There is already a body of literature around using these techniques in computational organic chemistry,^42^ and we believe that there are many more problems involving biochemical networks, gene regulatory networks, protein structure modeling, and computational neuroscience where these graph convolutional architectures can also be applied.

While there are obvious clinical applications of a successful phenotype classifier, there are many questions that still need to be answered before these techniques are accurate enough for use in the clinic. For example, we need to understand why the model makes certain predictions, because a medical practitioner will have more confidence in a prediction if it comes with an explanation. Recently, a class of algorithms called attribution methods have emerged which can identify predictive features in the input on a per-sample basis.^43,44^ This could be useful from a personalized genomics standpoint, as attribution methods could help explain which bacteria are contributing to dysbiosis in an individual patient and potentially suggest probiotic interventions to alleviate the dysbiosis. We will perform attribution analysis for our models to consider such questions in a future study.

Lastly, one of the shortcomings of this work is that it couples multiple questions together, making it difficult to study each one in isolation. For example, in addition to comparing different models for disease classification, we also made numerous assumptions in the construction of our dataset in terms of the diseases we included and the available metadata. We have not considered how factors such as gender, age, or body site might affect or be useful for enhancing a classifier’s performance. Additionally, there are numerous pieces of metadata from individual studies that could be used when analyzing these metagenomes. Although the original R package for curatedMetagenomicData makes it easy to explore the data, constructing training and test sets for deep learning took substantial effort. To make this exercise easier for other researchers, we have made our code available on Github, which contains not only our models and training scripts, but also our code for downloading and pre-processing the metagenome samples. We believe that the results shown here will be improved upon in the future as more studies are added to curatedMetagenomicData and better models and training procedures emerge. We hope that the code we provide will help other researchers attack this problem and similar problems involving machine learning with metagenomics.

## 6. Acknowledgments

Saad Khan was supported by the Einstein Medical Scientist Training Program (2T32GM007288-45). Libusha Kelly is supported in part by a Peer Reviewed Cancer Research Program Career Development Award from the United States Department of Defense (CA171019).

## External Links

Source: github.com/kellylab/Metagenomic-Disease-Classification-With-Deep-Learning

